# A foundation model for generalizable cancer diagnosis and survival prediction from histopathological images

**DOI:** 10.1101/2024.05.16.594499

**Authors:** Zhaochang Yang, Ting Wei, Ying Liang, Xin Yuan, Ruitian Gao, Yujia Xia, Jie Zhou, Yue Zhang, Zhangsheng Yu

**Affiliations:** Department of Bioinformatics and Biostatistics, School of Life Sciences and Biotechnology, Shanghai Jiao Tong University, Shanghai, China; Clinical Research Institute, Shanghai Jiao Tong University School of Medicine, Shanghai, China; SJTU-Yale Joint Center for Biostatistics and Data Science, Shanghai Jiao Tong University, Shanghai 200240, China; School of Mathematical sciences, Shanghai Jiao Tong University, shanghai, China

## Abstract

Computational pathology, utilizing whole slide image (WSI) for pathological diagnosis, has advanced the development of intelligent healthcare. However, the scarcity of annotated data and histological differences hinder the general application of existing methods. Extensive histopathological data and the robustness of self-supervised models in small-scale data demonstrate promising prospects for developing foundation pathology models. Due to the need for deployment, lightweight foundation models also need to be developed. In this work, we propose the BEPH (**BE**iT-based model **P**re-training on **H**istopathological images), a general lightweight foundation model that leverages self-supervised learning to learn meaningful representations from 11 million unlabeled histopathological images. These representations are then efficiently adapted to various tasks, including 2 cancer patch-level recognition tasks, 3 cancer WSI-level classification tasks, and 6 cancer subtypes survival prediction tasks. Experimental results demonstrate that our model consistently outperforms several comparative models with similar parameters, even with limited training data reduced to 50%. Especially when the downstream structure is the same, the model can improve ResNet and DINO by up to a maximum increase of 8.8% and 7.2% (WSI level classification), and 6.44% and 3.28% on average (survival prediction), respectively. Therefore, BEPH offers a universal solution to enhance model performance, reduce the burden of expert annotations, and enable widespread clinical applications of artificial intelligence. The code and models can be obtained at https://github.com/Zhcyoung/BEPH. And currently, online fine-tuning of WSI classification tasks is available for use on http://yulab-sjtu.natapp1.cc/BEPH.

## Introduction

Digital and computing tools can reduce the cost of the current healthcare system without compromising care standards, and may even improve its coverage and quality. In computational pathology, histopathological image analysis has been considered the gold standard for diagnosis.^1,2^. However, the conventional manual approach for pathology image diagnosis is time-consuming and demanding for pathologists. They need to carefully examine morphological features, such as tissue shape and cell sizes, throughout the entire image to provide accurate diagnosis. Insufficient diagnostic experience can lead to missed diagnoses and misdiagnoses, significantly impacting subsequent treatment^3^.

Advancements in computational pathology and artificial intelligence offer possibilities for objective diagnosis, prognosis, and treatment response prediction using gigapixel slides^4^. Particularly, deep learning-based computational pathology has shown promising prospects in various pathological tasks, including disease grading, cancer classification, and survival prediction^2,5–7^. Despite the revolutionary impact of deep learning on medical imaging, challenges remain to be addressed.

The typical approach for deep learning analysis of pathological images usually entails loading ImageNet pre-trained weights, and training the model using slides and specific labels. However, the inherent differences between natural scenes and pathological images affect the performance^8–10^. Researchers have recognized the positive impact of pre-training on pathological images, but their methods often involve supervised pre-training tailored to the specific problem. This approach is limited by the scarcity of training data and the lack of effective transfer of pre-trained weights to other tasks. Additionally, significant variations in morphology among different cancer types hinder the adaptation of pre-trained models to cross-cancer types. For more tasks, researchers need to continuously annotate data and develop new methods, reducing the efficiency of practical applications^11,12^. A promising solution to address these challenges is to pre-train foundation models using a wide variety and a massive amount of pathological images^13–15^.

Foundation models in computational pathology are developed by pre-training on a large volume of unsupervised digital pathology images. These models learn underlying structures and relationships in images, establishing a robust knowledge base. Subsequently, the foundation models are well fine-tuned using small labeled datasets customized for specific tasks and always achieve high- performance^16,17^. There are two main training strategies: supervised learning (SL) and self-supervised learning (SSL). SL involves pre-training on labeled data, which is challenging due to the large gap between the speed of data annotation and data acquisition18. In contrast, SSL utilizes pretext to extract information from large-scale unsupervised data, reducing reliance on labels. SSL methods include contrastive learning and MIM (Mask Image Modeling). Contrastive learning focuses on constructing positive and negative sample pairs ^19,20,22^, while negative samples are usually obtained from other images in the dataset or through self-transformation. If appropriate pairs of positive and negative samples are constructed, contrastive learning can optimize representations independently of downstream tasks and preserve meta-information to the greatest extent possible. However, constructing sample pairs in histopathological datasets can be challenging due to a strong resemblance between different images. Chen et al. proposed a contrastive learning pre-trained DINO framework, named HIPT, which achieved excellent performance on subtype classification and survival prediction^21^. Due to DINO’s bottom-up aggregation to learn image representation, this approach requires computationally costly hierarchical pre-training. Moreover, as they mentioned in the original paper, it is not simple to obtain general and discriminative representations of WSI directly^21^.

Although some studies have addressed the challenges of contrastive learning in pre-training pathological images^17,23,24^, the effectiveness of masked image modeling methods, another significant branch of visual models, has not been demonstrated in pre-training pathological images. At the same time, these studies focus more on the breadth of downstream tasks, with slightly insufficient analysis of depth. For instance, they do not include survival prediction, an essential clinical task, and fail to explain why self-supervised methods outperform other methods. This study aims to address these gaps by proposing a pre-training MIM foundation model for cancer detection and survival prediction. The model aims to achieve efficient prediction of various types of cancers, utilize heatmaps and Kaplan-Meier survival curve analysis^25^, to delve into the MIM, contrastive learning, and natural image supervised training performance differences, to enhance the interpretability of results. Besides, Large models typically require more computing resources for training and inference. For many researchers and some real-time applications or low-latency scenarios, the foundation model may be more suitable. Therefore, lightweight foundation models may make computational pathology more widely applicable in clinical applications.

In this work, we propose a novel lightweight SSL-based foundation model called BEPH. The parameter size is only 86 M, which can be deployed in computers easily. We systematically evaluate its performance and versatility in various cancer detection tasks. Fig. 1 provides an overview of the construction and application of BEPH. The histopathological images used for pre-training are sourced from the Cancer Genome Atlas (TCGA), encompassing 32 different types of cancer^26^ (Fig. 1b). We have collected a total of 11,760 pathology images (with 92 exclusions due to indeterminate magnification), resulting in a pre-training patch dataset of 11,774,353 samples. This dataset is 10 times larger than the ImageNet-1K dataset (1.28 million). Detail statistics for each cancer slides and patches are presented in Fig. 1b.

**Fig. 1.**
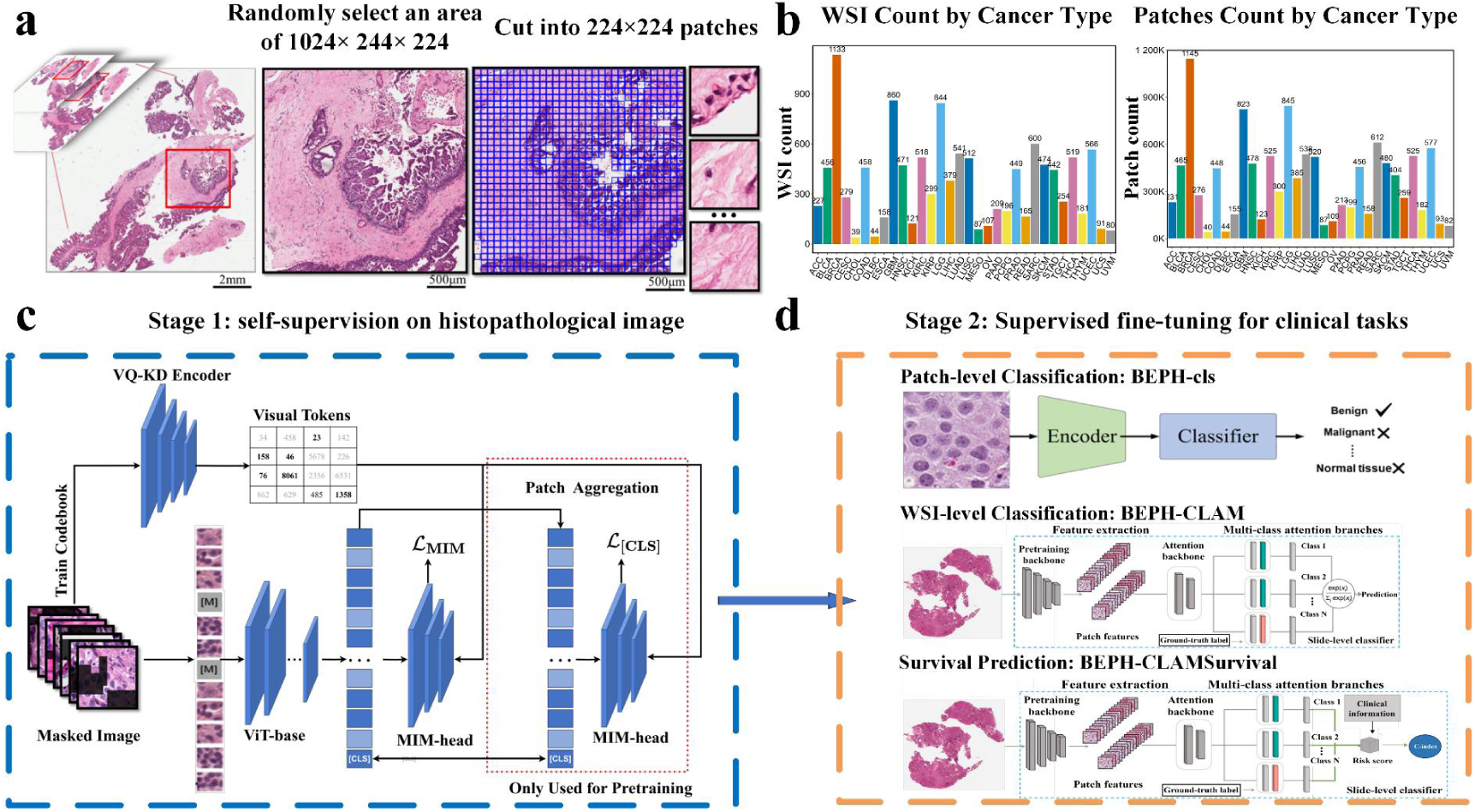
Overview of the BEPH architecture. **a**, The segmentation of local tissue regions from the WSI and the division into patches. **b**, Distributions of WSI and patches in the collected pre-training dataset. **c**, The BEPH architecture for pre-training. The pre-training phase employs the MIM method for SSL of pathological image features. **d**, The downstream supervised fine-tuning architectures, including three tasks: patch-level classification structure (BEPH-cls), WSI-level weakly supervised classification structure (BEPH-CLAM), and WSI-level survival prediction structure (BEPH- CLAMSurvival).

To efficiently train BEPH, we utilize advanced SSL techniques (specifically BEiT^27,28^) for pre-training on both natural images from ImageNet-1k and the collected pathology images. This involves initializing the network with pre-trained weights from natural images and then further pre-training on the TCGA to learn a generalized representation of pathology images^15,29^.

Subsequently, we fine-tune BEPH on a wide range of challenging classification and prediction tasks. We first consider the classic patch-level classification tasks. The downstream structure is BEPH-cls (Fig. 1d). Secondly, we explore the detection of WSI levels, which provide comprehensive and accurate pathological information. This work focuses on subtypes classification of BRCA, NSCLC and RCC (see Supplementary Table 1 for full name of cancer), called BEPH-CLAM (Fig. 1d). Finally, we considered survival prediction tasks in BRCA, CRC, CCRCC, PRCC, LUAD and STAD, named BEPH-CLAMSurvival (Fig. 1d). The data of BEPH-CLAM and BEPH-CLAMSurvival are from TCGA. Hence, our evaluation of the universality of cancer analysis is based on multi-task and multi-cancer (for specific details, please refer to the Methods). Compared to the state-of-the-arts models, including traditional transfer learning models, weakly supervised models, and contrastive learning models (DINO), BEPH consistently achieves superior performance and label efficiency in these tasks. Furthermore, we delve into the interpretation of BEPH’s disease detection performance through qualitative results and variable control experiments, which indicate that significant image regions reflect established knowledge from pathology and pathological literature. Finally, consider that foundation models like UNI and CONCH only allow researchers to perform inference using pre-trained base models. We has open-sourced our pre-training, fine-tuning, and inference code and deployed BEPH to http://yulab-sjtu.natapp1.cc/BEPH , serving as a foundation for various downstream tasks.

## Results

### Patch-level classification

We used the BreakHis^30^ dataset for binary classification of benign and tumor in cancer through five random experiments^31,32^. Unlike the image splitting strategy used in current top-performing algorithms, we downscaled the images by a factor of 3.125 to 224x224 pixels, resulting in a loss of a significant amount of image details. Even though, at the patient level and image level, BEPH achieved average ACC ± s.d of 94.05 ± 0.9928 and 93.65 ± 0.6730, respectively, which is 5%-10% higher than the latest reported CNN models and weakly supervised models. It also outperformed the best-reported performance of self-supervised models by 1.9% and 1.6% (Fig. 2a, b). Besides, BEPH exhibits consistently higher performance at certainly magnification levels at the image level and patient level compared to other algorithms (Fig. 2c, d).

**Fig. 2.**
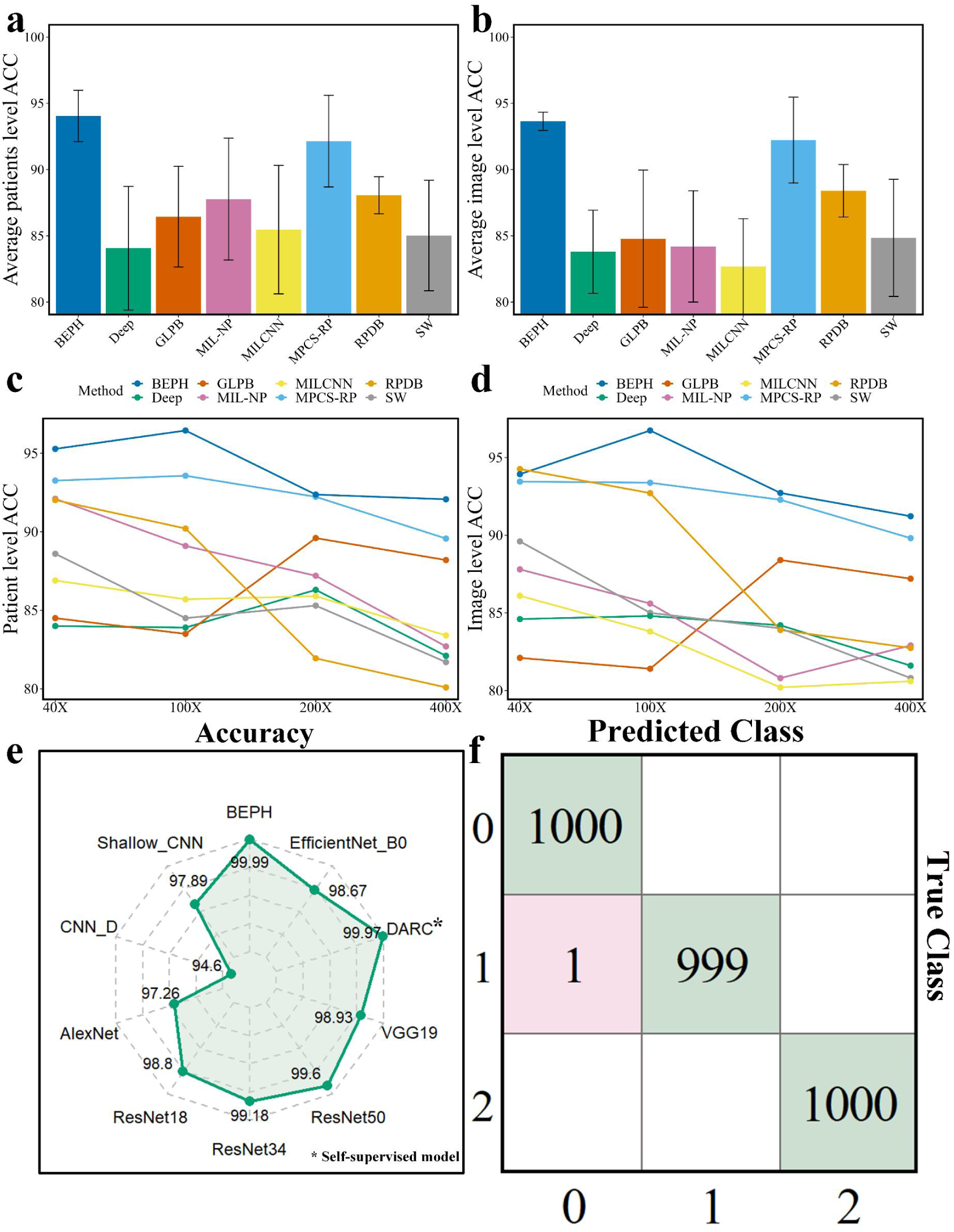
Performance evaluation on publicly available datasets composed of patches. **a**, Average accuracy comparison at the patient level for different resolutions. The compared models include CNN models (Deep^31^, SW^31^, GLPB^48^, RPDB^49^), weakly supervised models (MIL-NP^32^, MILCNN^32^), and self-supervised model (MPCS-RP^38^). **b**, Average accuracy comparison at the image level for different resolutions. The compared models are the same as a). **c**, **d**, Performance of different methods at patient level and image level, respectively, at magnification of 40, 100, 200, and 400. **e,** Performance of different algorithms on the LC25000 dataset. **f**, Confusion matrix on validation set of LC25000 dataset.

To test the generalization ability of the model across different cancers, we further deployed the model on LC25000 datasets^33^. BEPH achieved an average Accuracy score of 99.99% ± 0.03 across three lung cancer subtypes of 99.99%±0.03(Fig. 2f) which is higher than any other reported models like shallow-CNN^34^, AlexNet^35^, ResNet^35^, VGG19^35^ and EfficientNet-B0^36^ to self-supervised model DARC-ConvNet^37^ (Fig. 2e).

These results show that MIM-based pre-training enhances the adaptability of the model to pathological image resolution and cancer type transformation, and robustness to low-resolution image information, demonstrating the clinical potential of BEPH.

### WSI-level Classification

With the advancement of technology and improvement in computational power, modern research is increasingly focusing on the pathological information contained in whole-slide images (WSIs). Cancer subtyping provides a foundation for personalized treatment, helps predict patient prognosis, and contributes to understanding disease and mechanism research. Therefore, in this section, we train a weakly supervised subtype classification model based on a self-supervised feature extractor with multiple instance learning and aggregate attention (MIL). After randomly selecting a 10% independent test set, we evaluate the sliding-level WSI classification performance of three clinical diagnostic tasks using 10-fold cross-validation: RCC subtypes, NSCLC subtypes, and nonspecific invasive breast cancer.

For classification of the three subtypes of renal cell carcinoma (RCC): papillary renal cell carcinoma (PRCC), chromophobe renal cell carcinoma (CRCC), and clear cell renal cell carcinoma (CCRCC), using the public RCC WSI dataset, the model achieves a 10-fold macro-average test area under the receiver operating characteristic curve (AUC) of 0.994±0.0013 (Figure 3.a). For the two subtypes of non-small cell lung cancer (NSCLC) in the public NSCLC WSI dataset: lung adenocarcinoma (LUAD) and lung squamous cell carcinoma (LUSC), the model achieves an average test AUC of 0.970±0.0059 (Figure 3.d). In the public BRCA WSI dataset, for the two subtypes of nonspecific invasive breast cancer: invasive ductal carcinoma (IDC) and invasive lobular carcinoma (ILC), the model achieves an average test AUC of 0.946±0.019 (Figure 3.g). These results significantly outperform the corresponding models trained with pre-training on natural images or pathology images, such as MIL^2,3^ (DINO pretrained on 256×256 pathology images), GCN-MIL^3,4^ (DINO pretrained on 256×256 pathology images), DS-MIL^3,5^ (DINO pretrained on 256×256 pathology images), DeepAttnMISL^3,6^ (DINO pretrained on 256×256 pathology images), CLAM-SB ^4^ (ResNet supervised pretrained on ImageNet-1K), CLAM-SB ^4,3^ (DINO pretrained on 256×256 pathology images), and HIPT^3^ (DINO pretrained on 256×256 and 4096×4096 pathology images) (p-value < 0.05). Choosing DINO for comparison is because the parameter quantity is similar.

**Fig. 3.**
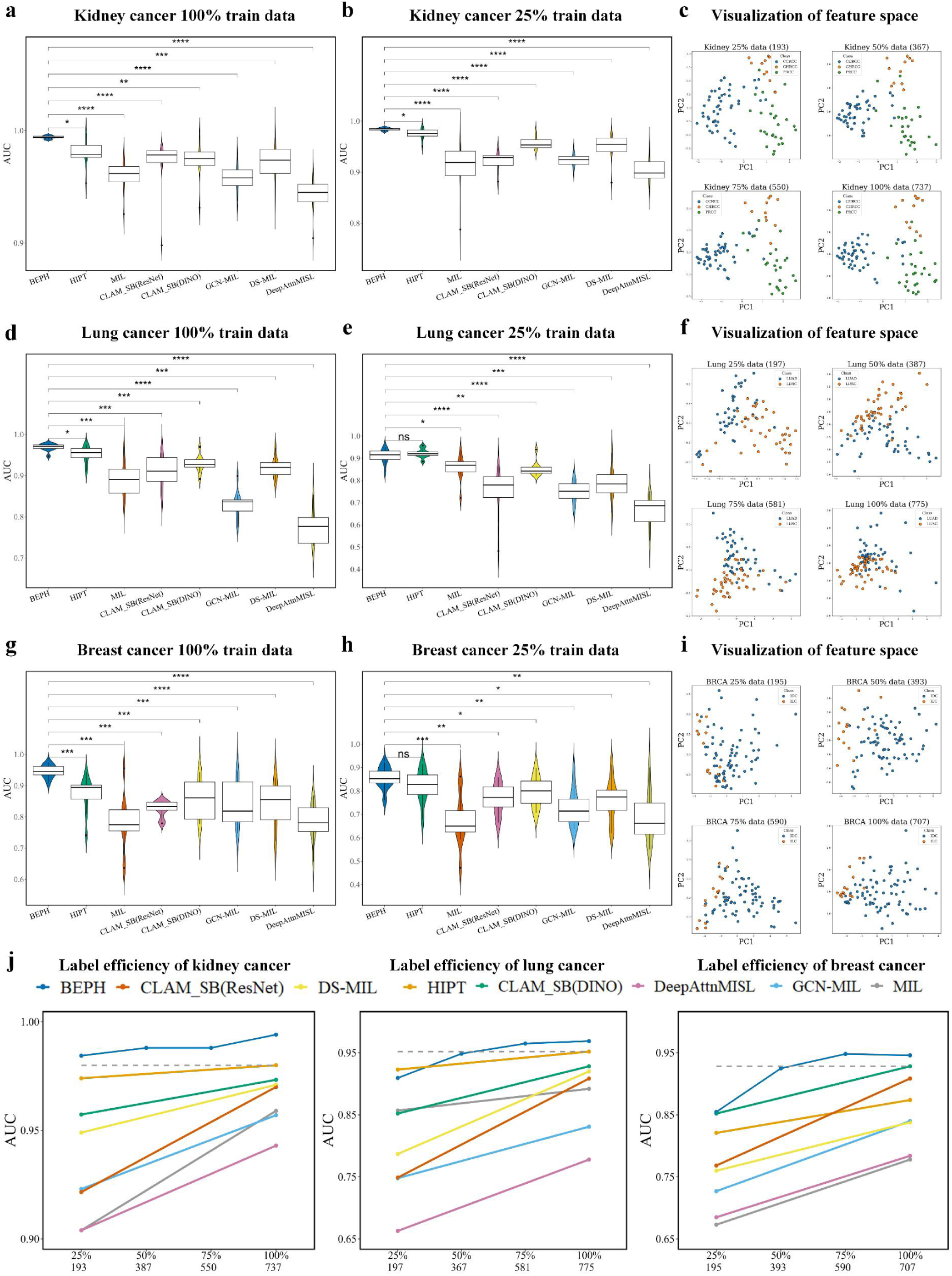
Weakly supervised classification results in the WSI-level independent test dataset. **a**, **d**, **g**, Independent test results for kidney cancer, lung cancer, and breast cancer using only 25% of the training data. **b**, **e**, **h,** Independent test results for breast cancer, kidney cancer, and lung cancer using 100% of the training data. **c**, **f**, **i**, Two-dimensional visualization of WSI-level feature space in BEPH, with different training data sizes. The number of slides used for each training data size is indicated in parentheses. With the increase of train dataset, the clusters are tighter, indicating that the true relationships between features are easier to capture and represent. **j**, Tolerance of different pre-training models to data reduction. The horizontal scale is the percentage and the corresponding number of samples. These results demonstrate that the MIM pre-training model can achieve good performance after training on a limited number of labeled slides, surpassing other weakly supervised baselines.

Specially, we selected the model for CLAM_SB to evaluate the impact of different pre-training methods on performance. The BEPH outperforms the ResNet supervised pre-trained on ImageNet- 1K (ResNet-ImageNet) by 8.8% in the BRCA, as well as outperforms the DINO pre-trained on 256×256 histopathological images (DINO-HistoPretrain) by 7.2%. There is a relatively small improvement in the kidney aspect, with an increase of 2.1% and 1.4%, which may be due to significant differences in features among PRCC, CRCC, and CCRCC subtypes. The ResNet- ImageNet already achieves good results (97.3%), but the performance can be further improved (99.4%) with MIM pre-training. This indicates that BEPH has learned valuable features and patterns from large-scale slides, enabling accurate classification and identification of pathological changes in cancer.

We also tested the zero-shot capability through cross-evaluation, to ensure its reliability and usability across different datasets and real-world scenarios. When fine-tuning the model on the CAMELYON16 dataset^41^, the BEPH achieved a score of 0.9056 ± 0.0221 in the internal validation set, which is comparable to the performance of the state-of-the-arts models (Supplementary Fig. 1). Furthermore, the AUC tested on the BACH dataset^42^ yielded a score of 0.638 ± 0.0585, which was statistically significantly higher by 20% than other models (P < 0.05). Among all the self-supervised and weakly supervised models, the zero-shot capability of BEPH is the most powerful. During cross-evaluation, the AUC results were noticeably lower than those obtained in the internal validation data, indicating a significant improvement in downstream task performance through fine-tuning.

### Online Fine-tuning in the WSI classification task

Although with the complete open source of code repositories such as GitHub, many researchers can easily access, understand, and use advanced foundational models. there are still significant barriers for researchers or doctors who have not been exposed to artificial intelligence to deploy and reproduce code. Therefore, in order to promote automated diagnosis of various cancers, we will deploy BEPH to http://yulab-sjtu.natapp1.cc/BEPH for use by all researchers. At present, researchers can complete online fine-tuning of WSI classification tasks by uploading WSI image data according to the tutorial.

### Label efficiency for cancer detection

The foundation models require good label efficiency, because acquiring labeled WSI data is challenging, especially for rare diseases. When the training data was reduced to 25% of the original size while keeping the test dataset unchanged, the results indicate that even with a significant reduction in the proportion of training data, the performance of the BEPH model remains superior to other weakly supervised models (Fig. 3b, e, h). Compared with HIPT, we observed a significant improvement in kidney cancer, and the classification AUC of BEPH in invasive breast cancer was 3.4% higher than that of HIPT. The limited sample size may account for the insignificant difference in SSL models. We also created different subsets of training data, including 25%, 50%, 75%, and 100% of the original data (Fig. 3j). It can be observed that when approximately 50% of the training data is used, the performance of BEPH is comparable to other models trained with 100% of the data. In the case of kidney cancer, even a reduction to 25% achieves the best performance. This indicates that MIM pre-training enables the model to better tolerate data scarcity issues.

### Survival Prediction

Clinical survival prediction models’ development contributes to improving clinicians’ ability to evaluate patients’ prognosis. Therefore, we trained a survival risk regression model based on a self-supervised feature extractor and weakly supervised learning. BEPH ranked first in predictive capabilities among all models in each of the six different cancer types: BRCA, CRC, CCRCC, PRCC, LUAD, and STAD. The C-index values were 0.6643 (95% CI 0.6234, 0.7052), 0.6758 (95% CI 0.5863, 0.7652), 0.6685 (95% CI 0.6206, 0.7164), 0.7135 (95% CI 0.5203, 0.9068), 0.6039 (95% CI 0.5093, 0.6986), and 0.5941 (95% CI 0.5348, 0.6534), respectively. Overall, the performance improvement in C-index ranged from 1.1% to 5.5%. In pairwise comparisons among different models (Supplementary Table 4) within each cancer dataset, BEPH achieved the highest C-index performance in all six cancer types (Fig. 4a). Among of them, the highest C-index is 0.7135 in PRCC. Similarly, we chose the CLAMSurvival model to evaluate the impact of different pre-training methods. The BEPH shows significant performance improvement (6.44% and 3.28% on average) compared with the ResNet-ImageNet, and the DINO-HistoPretrain. Further, we analyzed the ability of these methods to model prognostic risk groups ^43^. Fig. 4b shows that BEPH is superior in distinguishing between high- and low-risk patients, on the cancers mentioned above. For example, on CCRCC, BEPH recognizes high- and low-risk patients, with a p-value smaller than 0.0001 (vs. a p-value of 0.00071 given by DINO-HistoPretrain and a p-value of 0.0002 given by ResNet- ImageNet). On CRC, our method also brings a significant difference with a p-value of 0.016 (p-value < 0.05), while DINO-HistoPretrain and ResNet-ImageNet fails to distinguish high- and low-survival risks significantly (p-value = 0.21 and p-value = 0.51) (for other cancers, see Supplementary Fig.2).

**Fig. 4.**
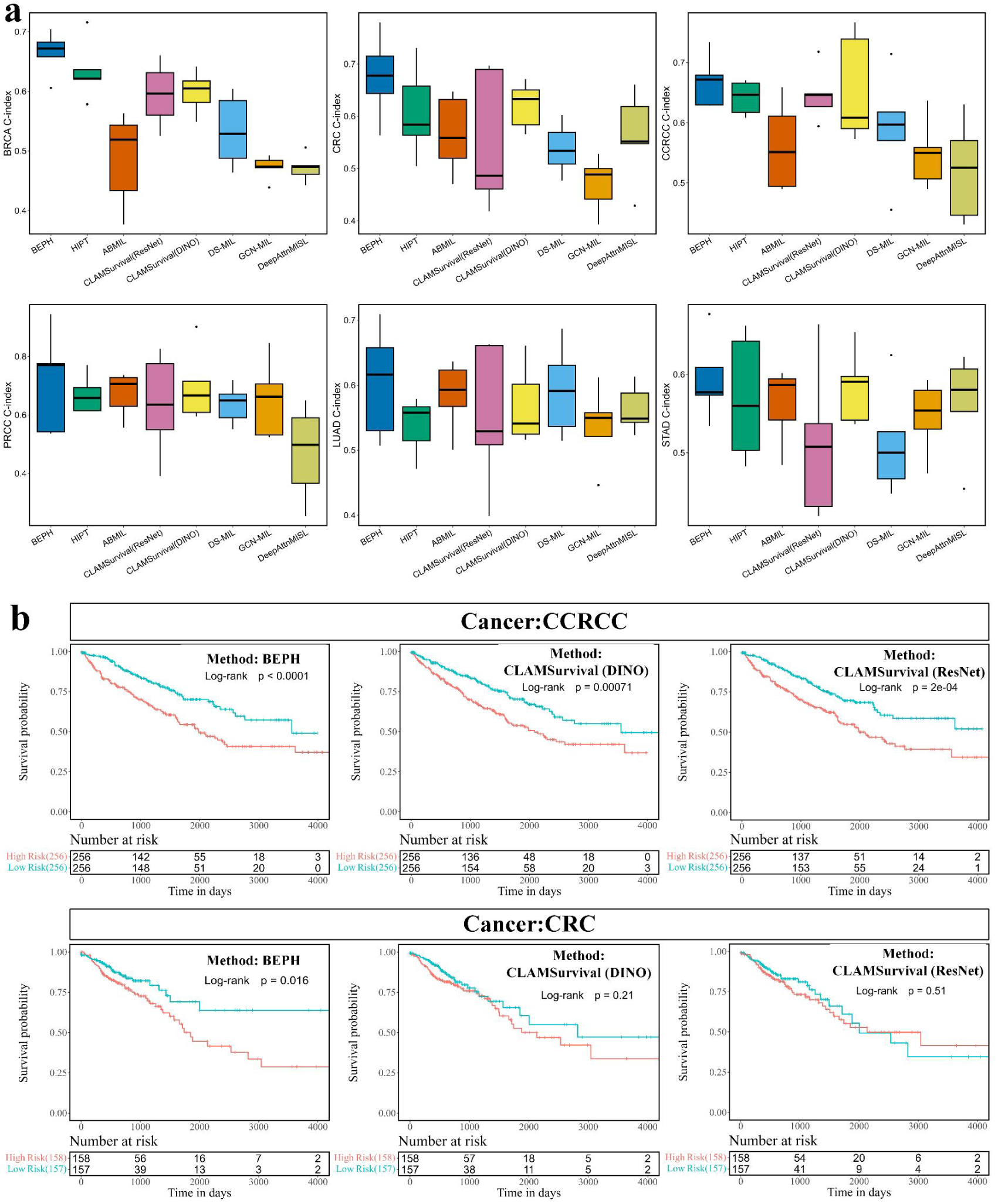
Survival Prediction. **a**, Different methods of C-index on six cancers. BEPH exhibits higher C-index compared to weakly supervised models and baseline models across six different cancer subtypes. **b**, Survival curves for patient stratification are depicted using Kaplan-Meier analysis. These curves illustrate the true survival rates of patients classified as high or low risk. The number of patients who survived in each risk group is shown below the survival curves. The log-rank test calculates the p-value, indicating the significance of the difference in survival between the two groups.

These indicate that BEPH is adept at identifying survival-related patterns and better predicts patients’ survival. Furthermore, it doesn’t require annotation on histopathological images or the introduction of additional information such as genomes, making it more practical for real-world applications.

### Whole-slide attention visualization and Interpretability

To gain a deeper understanding of the exceptional performance of BEPH in downstream tasks, we conducted a qualitative analysis of BEPH’s application in self-supervised pre-training and specific task decision-making. We studied the patch-level feature space learned by the BEPH model. We introduced an external dataset, NCT-CRC-HE-100K^44^, and used UMAP to reduce its instance-level 768-dimensional feature representation to two dimensions (only using the backbone for feature extraction without involving downstream tasks). It can be observed that patches belonging to different prediction categories were clustered into distinct clusters in the feature space. The performance of feature extraction pre-trained on histopathology images is superior to those solely pre-trained on ImageNet-1k, and significantly better than randomly initialized parameters. This is particularly evident in the smooth muscle (MUS) and debris (DEB) categories (Fig. 5a).

**Fig. 5.**
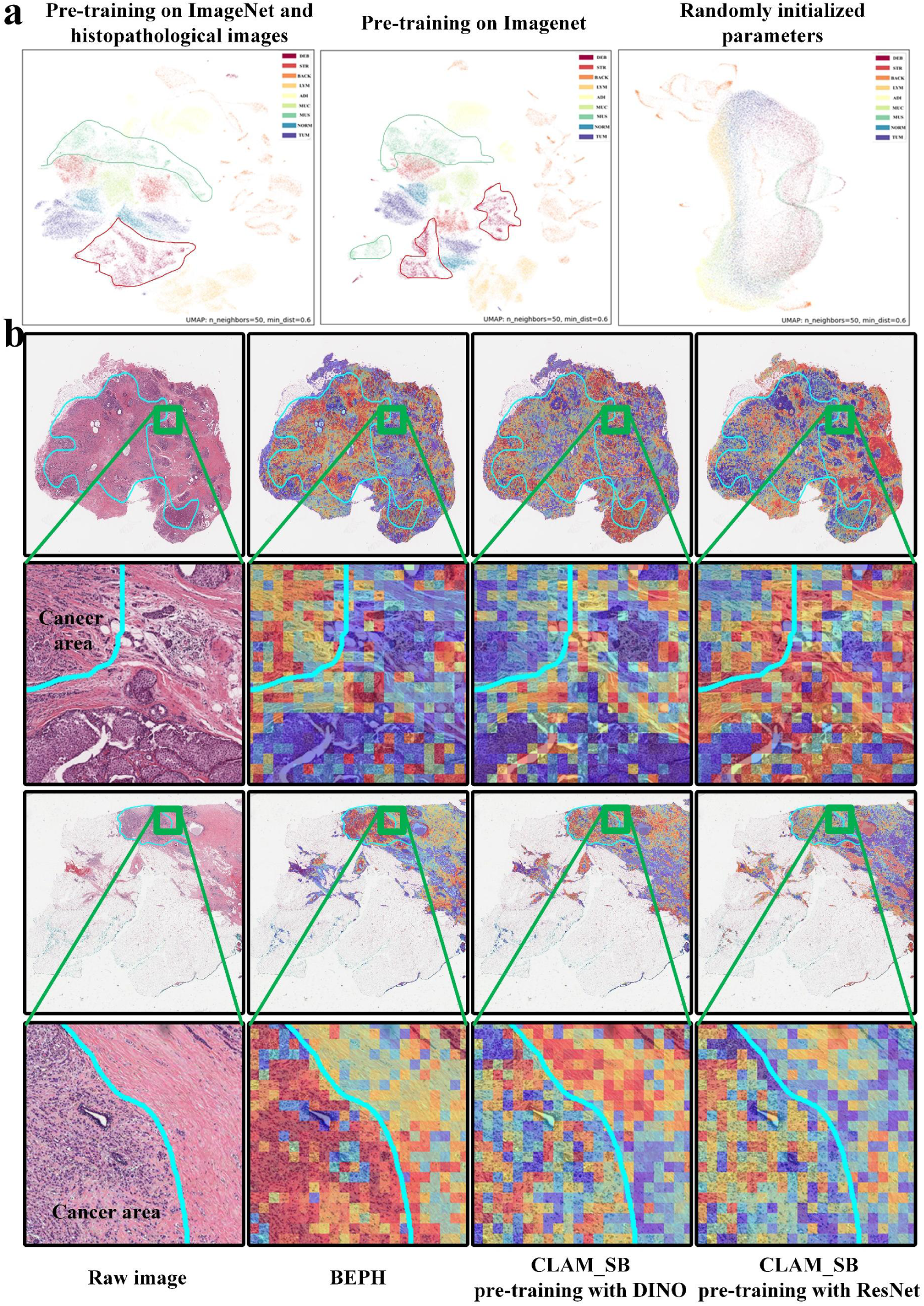
BEPH Model Interpretability and Visualization. **a**, The interpretability and visualization of the pre-trained model are demonstrated after extracting features from the NCT-CRC-HE-100K dataset and applying UMAP dimensionality reduction. The closer the instances of the same class are, the better the model’s feature extraction capability. **b**, The figure displays automatically calculated attention heatmaps covering WSI. The warm colors in the heatmap indicate higher estimated probabilities of tumor tissue. The or blue contour lines represent cancer regions annotated by pathologists, and the green boxes denote local areas containing the boundary between tumor and normal tissue. From left to right, the first column shows the ground truth, the second column displays BEPH, the third column shows DINO-HistoPretrain, and the fourth column presents ResNet- ImageNet. The odd rows depict the model’s attention on the entire tumor region of the WSI, while the even rows show the heatmap for the corresponding local area within the box. These slides are from the TCGA-BRCA dataset.

Additionally, we visualized the WSI-level feature space using principal component analysis (PCA) reducing the feature vectors to two-dimensions (Fig. 3c, f, i). The results show that as the training data increases, the clusters of different categories in the sample space become tighter, and show clear boundaries between different categories, indicating that an increase in training data helps the model learn features better.

To interpret the fine-tuning performance, we employed attention scores to visualize the salient regions of the images in the downstream tasks. We found that even without explicitly informing the model about the presence of tumor tissue and its subtypes, just using weakly supervised learning with WSI-level labels, the attention heatmaps exhibited high consistency with the annotations of pathologists across all tasks. This indicates that the downstream fine-tuned model can accurately identify tumor and tissue regions within a limited number of iterations (20 epochs) (Fig. 5b). On the other hand, ResNet-ImageNet and DINO-HistoPretrain show noticeable biases in their attention towards tumors, which may explain their poor performance compared with BEPH (Fig. 5b).

### Ablation test

To investigate the effectiveness of pre-training using pathology images to enhance the ability of self-supervised models, we conducted ablation experiments. Fig. 5a demonstrates that the model pre-trained with pathological images exhibits superior feature extraction ability than that pre-trained solely with ImageNet-1k. To quantitatively evaluate the effect of the improvement, we performed patch-level evaluation on BreakHis, WSI-level evaluation on NSCLC subtype and survival prediction for BRCA, respectively. Table 1 shows that the performance of the model pre-training on pathological images is better than that of the model pre-trained only on ImageNet-1K (BEiTv2- ImageNet). This highlights the difficulty of SSL using natural images for specialized domains and emphasizes the importance of providing a large amount of pre-training data from the specific field which helps to gain a deeper understanding and enhance the generalization ability. Complete ablation test results are in Supplementary Tables.

**Table 1.** Ablation experiments on the effectiveness of pre-training.

|  | **BreakHis (Mean resolution)** |  | **WSI weakly Classification** |  | **Survival prediction** |
| --- | --- | --- | --- | --- | --- |
|  | Patient Level | Image Level | NSCLC | NSCLC | BRCA |
|  | Accuracy | Accuracy | 25% | 100% |  |
| BEiT<sub>v2</sub>-ImageNet | 92.60±2.324 | 92.25±1.888 | 0.8788±0.021 | 0.9270±0.013 | 0.6440±0.0354 |
| BEPH | **94.05±0.9928** | **93.65±0.6730** | **0.9095±0.0332** | **0.9719 ±0.0054** | **0.6643±0.0330** |

## Discussion

Altogether, we utilized the MIM approach to create a lightweight foundation model for tissue pathology, named BEPH, in large-scale unlabeled pathological images. The contributions of this work are as follows. Firstly, we developed a self-supervised model for 32 cancer histopathological features from TCGA. The key technical advantage is utilizing SSL to learn features from histopathological images. By Training a histology-specific feature extractor on 11 million unlabeled patches demonstrates the effectiveness of this approach. Secondly, BEPH adapts effectively to various disease detection tasks by learning image representations through the pretext task of reconstructing images from heavily masked patches. It improves patch-level and WSI cancer diagnosis, as well as survival risk prediction. Comparative analysis with ImageNet pre-trained models and contrastive self-supervised models highlights the importance of MIM pre-training on large-scale datasets, even with limited training data. Additionally, after limited downstream task training, BEPH generates localized tumor heatmaps that match pathologists’ annotations. Combined with two-dimensional feature scatter plots, it is shown that MIM pre-training using tissue images enhances the model’s attention to specific tissue features.

The MIM model in this study is based on the BEiTv2 architecture for three reasons. Firstly, BEiT pioneered MIM method, and BEiTv2 outperforms models like MAE and CAE^45^ comprehensively. Secondly, both BEiT and BEiTv2 use discrete visual tokens, which are more suitable for classification tasks. Models like MAE struggle to learn features at multiple scales and finer granularity, resulting in uniform features from pathological maps and limited exploration of deep semantic information. However, MAE has stronger interpretability due to its ability to reconstruct RGB pixels. Therefore, a potential research direction could involve augmenting the MAE model structure to learn multi-scale and fine-grained features and pre-training it on pathological images. BEPH has broad applications in cancer detection tasks. When pixel annotations are available to segment cancerous tissues, BEPH can perform patch-level classification tasks. Even without pixel annotations, the model can take the entire histopathological image as input and perform weakly supervised learning for WSI classification tasks. In cases where clinical information is available, BEPH can be transformed into a survival prediction model to predict patients’ outcomes. However, it is important to note that various indications suggest that fully supervised patch-level classification outperforms weakly supervised WSI-level classification. Therefore, it is recommended to collect pixel annotations whenever possible in practical applications.

The use of CLAM as a WSI-level classification model is based on three considerations. First, since only a small portion of the image is relevant to its class label, performing supervised classification at the patch level would introduce significant bias by labeling all patches as the same class. Second, CLAM, as an attention-based MIL model, handles imprecise annotations effectively. Its plug-and-play nature allows convenient application of pre-training results. Third, BEiT, similar to large language models, typically possesses contextual learning capabilities, which can compensate for CLAM’s limitation of failing to learn potential nonlinear interactions between instances. This may be one of the reasons why the weakly supervised classification performance of CLAM pre-trained with MIM is superior to that of CLAM pre-trained with DINO.

Our work represents an important step towards establishing a foundation pathology model. However, there are still some issues that need to be addressed. Firstly, although our self-supervised pre-training dataset is much larger compared to currently reported datasets, it is limited to TCGA and fails to effectively utilize all pathological tissues. Secondly, due to limited computing resources and the need for fine-tuning on web servers, we have chosen a more direct ViT-base backbone instead of a ViT-large. However, it is worth noting that with more parameters, ViT-large may potentially enhance the model’s representational capacity, which could be a direction for future improvements. Additionally, as the MIM model continues to evolve, the pre-trained models currently used may limit the performance ceiling of future foundation models. Thirdly, even the training time and parameters are consistent with other big vision models of similar scale and the loss reaches a convergent state, we still don’t have sufficient evidence that BEPH has fully learned the features of pathological images. In theory, longer training time generally leads to better performance. It remains uncertain whether further extending the training time would improve the performance.

In the future, our plans involve constructing larger-scale pre-training datasets that include a greater variety of pathological images from different sources, allowing the pre-trained models to be more extensively. Additionally, we aim to explore more efficient methods for foundation model pre-training. One approach is to combine the advantages of contrastive learning and MIM, like the CMAE (Contrastive Masked Autoencoders) model, which may further enhance the capabilities. Moreover, it is necessary to investigate other popular computational pathology tasks, such as nucleus and gland detection and segmentation. Furthermore, the potential path towards achieving general artificial intelligence in pathology lies in multimodal large models that incorporate data from multiple domains like omics, text, and images. Exploring the integration of multimodal data holds significant research value in the field.

## METHODS

### BEPH architecture and implementation

The BEPH architecture is based on a default configuration of BEiTv2, which consists of an autoencoder and an Encoder similar to BERT. The Encoder encodes the original images into visual tokens using an autoencoder, VQ-KD, and then randomly masks 40% of the tokens. Finally, the tiles are fed to a backbone--ViT-base. The pretext task for pre-training is to predict the visual tokens of the original image based on the masked image tiles. This part is similar to the idea of Masked Autoencoder (MAE), but instead of predicting normalized image details, it predicts the visual tokens generated by VQ-KD.

To address the discrepancy between patch-level pre-training and image-level representation aggregation, a Patch Aggregation structure is added during pre-training. This structure shares the information of [CLS] token and the parameters of the MIM (Mask Image Modeling) head to better learn global representations.

The overall loss function minimizes the difference between the predicted tokens and the ground truth tokens (Loss_[MIM]_) and the global loss (Loss_[CLS]_). The total loss is the sum of these two losses. In our experiments, the model parameter size is 86 M. The image size is 224**×**224 and the batch size is 1024. The optimizer is AdamW and the initial learning rate is 0.0015. Finally, our model is pre-trained for 340 hours

### Dataset processing details for pre-training

For each digitized slide, our process for pre-training WSI begins with the automatic segmentation of the tissue regions. The slide is loaded into memory at a downsampled resolution (e.g., 40×). After thresholding the saturation channel of the image, a binary mask of the tissue regions (foreground) is generated. Considering that some slides in the TCGA dataset may contain duplicated pathological tissue and the slide has a large size, it is unnecessary to include all the patches in pre-training. To preserve the contextual relationship between selected patches, we locally sample image regions of 1024×224×224 (approximately 1024 images) from each pathological image, ensuring that the sampled region has a tissue proportion greater than 75%. These sampled image regions are then cropped into 224×224 tiles, while maintaining a tissue proportion of 75% and above within each patch (Supplementary Fig.3). In the end, we obtain an unlabeled pre-training dataset consisting of 11,774,353 tiles. This pre-training dataset is 10 times larger than the ImageNet-1K dataset (1.28M).

### Adaptation to downstream tasks

#### Patches level classification--BEPT-cls

When adapting to downstream tasks, we only use the encoder (ViT-base) of the base model and do not require the decoder. For patch-level classification tasks, we connect a single layer perceptron to the output of the encoder. The input image is encoded into a 768-dimensional feature representation by the encoder and fed into the perceptron, which outputs the probabilities of cancer categories. The category with the highest probability is defined as the final classification. The number of categories determines the number of neurons in the last layer of the perceptron.

The training objective is to generate classification outputs that match the labels. The batch size is set to 128, and the total number of training epochs is 30. The first 20 epochs are used for learning rate warm-up (from 0.0001 to a learning rate of 0.005), followed by a cosine annealing schedule where the learning rate decreases from 0.005 to 1 × 10^-6^ for the remaining 10 epochs.

For BreakHis, the dataset was randomly divided into training sets (80% of cases) and validation sets (20% of cases) for five repeated trials. If patches were classified based on patients, they were proportionally allocated by patient, and the average results of the validation sets from the five trials were taken as the final result. For LC25000, the dataset was divided into 5 folds and the average results of the validation sets from the five folds were taken as the final result.

#### WSI weakly supervised classification—BEPH-CLAM

For weakly supervised classification tasks with WSI, we freeze the encoder (ViT-base) and only use it as a feature extractor. The downstream structure is based on an attention-based Multiple Instance Learning (MIL) framework similar to CLAM. The encoder generates high-level features from the pathological images, and the attention network aggregates patch-level information into WSI-level representations for making final diagnostic predictions.

All models are trained for 20 epochs with early stopping. Early stopping is triggered when the validation AUC does not improve from the previous highest point for consecutive 10 epochs. The learning rate is adjusted slightly based on the cancer type and the proportion of training data, typically ranging from 2e-4 to 8e-4. Other parameters remain consistent with CLAM.

In the subtyping classification task, we reserve 10% of each dataset as test dataset and the remaining 90% for 10 fold monte carlo cross validation. In each fold, the model’s performance on the validation set was monitored during training and used for model selection, while the test set was kept separate and referenced only once after training for model evaluation. The dataset partitioning was slightly different from CLAM because CLAM fixed the validation set. To examine the model’s performance on training data, both the training and validation sets should be considered as part of the training data. Although reducing the validation set may affect model selection, it can test the model’s stability. All data for training, validation, and testing were performed at a 20**×** magnification.

#### WSI survival Prediction—BEPH-CLAM Survival

For the survival prediction task, we improved the final output layer of CLAM based on the survival model proposed by Wei et al., and adopted the COX proportional risk model to meet the requirements of survival prediction^46,47^, named BEPH-CLAMSurvival (Fig. 1d). The implementation is similar to the weakly supervised classification task, but with an additional linear layer in the last MIL layer that outputs a risk score, transforming it into a survival regression task. It’s important to note that the MIL framework is modified to incorporate clinical information.

The spatial features of the histopathological images are extracted by the encoder. The MIL framework is then utilized to estimate the risk function *ĥ*_θ_(*x_i_*), where θ represents the network’s weights. The weights of the network are trained based on the occurrence time (death) events to optimize the Cox likelihood function.

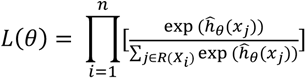

When we minimize negative log partial likelihood, the loss function is:

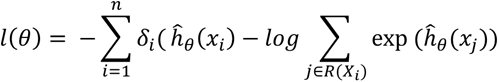

*δ_i_* is an indicator of whether the survival time is censored (*δ_i_* =0) or observed (*δ_i_*=1). θ is the weight of the CNN. ℛ(*X_i_*) denotes the set of individuals who are at risk for failure time of individual *i*.

The data was divided using a 5-fold cross-validation method, and both training and validation were performed at a 20× magnification level. The model undergoes a maximum of 20 epochs of training, and early stopping is implemented to prevent overfitting. Early stopping is triggered when the validation loss does not decrease from the previous lowest point for consecutive 5 epochs. The learning rate is adjusted slightly depending on the cancer type, ranging from 2e-4 to 2e-3. This range allows the model to find an optimal learning rate for each specific cancer type while training on the survival risk prediction task with WSI. Other parameters remain consistent with CLAM, ensuring consistency in the overall approach and framework utilized for the task.

#### Prognostic risk grouping

Patients are categorized into either a high-risk prognostic group (short-term survivors) or a low-risk group (long-term survivors) based on their hazard predictions compared to a predefined cutoff value. This cutoff value is determined by the median of hazard predictions. To evaluate the various pre-training methods, we employ CLAMSurvival on six cancer datasets and compare the performance of BEPH, DINO (contrastive learning pre-training), and ResNet (natural image supervised pre-training). The log-rank test is utilized to calculate p-values, which assess the differences in survival between the risk groups. It is important to note that the patients’ hazard predictions included in the risk grouping analysis are from the validation set of the 5 fold cross validation.

#### Evaluation Metrics for Fine-tuning

In patch-level classification tasks, accuracy is used to measure performance, which includes two scales: image-level accuracy and patient level accuracy (if applicable). Patient level accuracy reflects the performance of classification at the patient level and is calculated as the average of all patient scores. The patient score represents the proportion of correctly classified images within a patient’s set of images. On the other hand, image-level accuracy does not consider patient-level details. It is calculated as the proportion of correctly classified images among all images, regardless of the patients they belong to^38^.

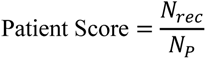

Let *N_p_* represents the number of pathological images owned by patient p. For each patient, *N_rec_* is the number of pathological images that are correctly classified.

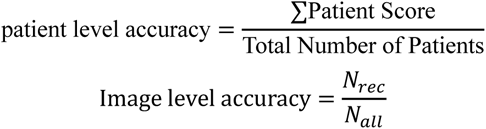

*N_all_* represents the total number of pathological images from all patients, and *N_rec_* represents the number of correctly classified pathological images among them.

In the classification task of WSI, the performance is evaluated using the AUC (Area Under the Receiver Operating Characteristic) metric. This metric is derived from the receiver operating characteristic curve of the classifier. For the BRCA and NSCLC subtype classification tasks, AUROC is calculated in a binary setting. For RCC subtype classification, AUC is calculated for each disease category separately, and then the average AUC is obtained. For survival prediction, the Concordance Index (C-Index) is a frequently used evaluation metric. Its primary purpose is to assess the survival models’ capacity to rank individual survival risks, where a higher prediction of risk is anticipated if the patient succumbs earlier. It can be written by

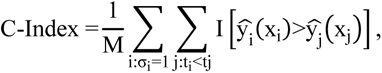

where M is the number of comparable pairs and I[·] is the indicator function. It ranges from 0.0 to 1.0. The larger the C-Index is, the better the model performance could be. The 95% confidence interval (CI) is obtained by 1.96×standard error. All significance tests in this paper are based on Wilcoxon test.

### Explanations for fine-tuned models

In the WSI classification task, the heatmap is similar to CLAM. we divide the slides into overlapping patches of size 224×224 and calculate the original attention scores for each patch. Then, we convert the attention scores into percentile scores, scaling them between 0 and 1.0. Afterward, we use a diverging colormap to convert the normalized scores into RGB colors. These colors are then displayed on the spatial locations of the slide, allowing for the intuitive identification and interpretation of high attention areas, which are shown in red to indicate areas of high attention.

### Datasets

#### BreaKHis dataset

The BreakHis dataset consists of 2480 histopathological microscope images of benign tissue and 5429 images of malignant tissue from 82 patients. The images were captured at magnifications of 40×, 100×, 200×, and 400×. Each image in the BreakHis dataset has a size of 700×460 pixels and was stained using HE staining technique. In the experiment, we compressed the dataset to 224×224 and grouped it by patient.

#### LC25000 dataset

The LC25000 dataset comprises 25,000 color images divided into 5 categories, with each category containing 5,000 images. All images have a size of 768 × 768 pixels and are stored in the JPEG file format. The dataset consists of the following 5 classes: colon adenocarcinoma, benign colon tissue, lung adenocarcinoma, lung squamous cell carcinoma, and benign lung tissue. 15000 images of lung tissues (5000 images of benign lung tissue,5000 images of lung squamous cell carcinoma and 5000 images of lung adenocarcinoma) were used in the experiment.

#### CAMELYON16 dataset

The CAMELYON16 dataset is a dataset for the detection of breast cancer with lymph node metastasis. It consists of 270 annotated whole slides for training and another 129 slides as a held-out official test set collected at the Radboud University Medical Center and the University Medical Center Utrecht in the Netherlands.

#### BACH dataset

The dataset comprises Hematoxylin and Eosin (H&E) stained breast histology whole-slide images in four classes: normal, benign, in situ carcinoma, and invasive carcinoma. We divide them into two classes: normal, tumor. The dataset contains 20 unlabeled normal, 10 pixel-wise labeled and 20 non-labeled slides. We only used normal and labeled slides.

#### NCT-CRC-HE-100K dataset

The NCT-CRC-HE-100K dataset is a collection of 100,000 non-overlapping image patches extracted from 86 human cancer tissue slices and normal tissue. The tissue slices were stained with Hematoxylin and Eosin (H&E). The dataset is sourced from the NCT Biobank (National Center for Tumor Diseases) and UMM Pathology Archive (Mannheim University Medical Center). Pathologists manually annotated the tissue regions in the whole slide images into nine tissue categories, which include: adipose tissue (ADI), background (BACK), debris (DEB), lymphocytes (LYM), mucus (MUC), smooth muscle (MUS), normal colon mucosa (NORM), stroma related to cancer (STR), and colorectal adenocarcinoma epithelium (TUM).

### Subtype classification dataset

#### The public BRCA WSI dataset

Our public BRCA dataset consists of 875 diagnostic WSI from the TCGA-BRCA database. Among them, 726 slides are from IDC (Invasive Ductal Carcinoma) and come from 694 cases, and 149 slides are from ILC (Invasive Lobular Carcinoma) and come from 143 cases.

#### The public NSCLC WSI dataset

Our public NSCLC dataset consists of 958 diagnostic Whole Slide Images from the TCGA NSCLC repository under the TCGA-LUSC and TCGA-LUAD projects. It includes 492 LUAD (Lung Adenocarcinoma) slides from 430 cases and 466 LUSC (Lung Squamous Cell Carcinoma) slides from 432 cases.

#### The public RCC WSI dataset

Our public RCC dataset consists of 903 diagnostic whole slide Images from the TCGA RCC database, including TCGA-KICH (Kidney Chromophobe), TCGA-KIRC (Kidney Clear Cell Renal Carcinoma), and TCGA-KIRP (Kidney Papillary Renal Cell Carcinoma) projects. It includes 118 CHRCC (Chromophobe Renal Cell Carcinoma) slides from 107 cases, 497 CCRCC (Clear Cell Renal Cell Carcinoma) slides from 491 cases, and 288 PRCC (Papillary Renal Cell Carcinoma) slides from 266 cases.

### The public survival prediction WSI dataset

The subtype dataset includes IDC and ILC pathology images and clinical information from the TCGA-BRCA database, comprising 726 images from 694 patients. The CRC (Colorectal Cancer) subtype dataset includes 315 images from 315 patients, sourced from the TCGA-READ and TCGA-COAD databases. The CCRCC subtype dataset is sourced from the TCGA-KIRC database and consists of 497 images from 491 patients, while the PRCC subtype dataset is sourced from the TCGA-KIRP database and consists of 288 images from 266 patients. The LUAD dataset is a subtype of lung cancer, sourced from the TCGA-LUAD project, and consists of 492 images from 430 patients. The STAD subtype dataset is one of the gastric cancer subtypes, including 348 pathology images from 348 patients, sourced from the TCGA-STAD database.

It’s important to note that the size of our dataset heavily references HIPT and CLAM, so the number of WSI in different datasets may not align exactly with the latest TCGA database.

### Computing hardware and software

SSL Model typically requires a powerful GPU for large-scale training and learning. We used 8 NVIDIA HGX A100 (40 GB) on the cloud platform of Shang hai Jiao Tong University. Downstream fine-tuning is also performed on NVIDIA HGX A100 (40 GB), with an average of 60 minutes required for every 500 WSI calculations.

All charts are generated using R (4.2.1). The AUC of the receptor operation characteristic curve was estimated using the Mann Whitney U statistical method, and its algorithm implementation was provided by the scikit learn scientific computing library (version 0.22.1).

## Data availability

TCGA diagnostic pathology images data (BRCA, NSCLC, RCC, CRC, CCRCC, PRCC, LUAD, STAD) and corresponding clinical data can be obtained from the National Cancer Institute’s Genomic Data Commons (https://portal.gdc.cancer.gov). CPTAC whole-slide data (NSCLC) and corresponding labels can be obtained from the National Cancer Institute’s Cancer Imaging Archive (https://cancerimagingarchive.net/datascope/cptac). BreakHis can be publicly accessed at https://web.inf.ufpr.br/vri/databases/breast-cancer-histopathological-database-breakhis/. The LC25000 dataset can be obtained from here: https://academictorrents.com/details/7a638ed187a6180fd6e464b3666a6ea0499af4af. NCT-CRC-HE-100K can be obtained from https://zenodo.org/records/1214456.

## Code availability

All code is implemented in Python, using PyTorch as the main deep learning library. The pre-training framework relies on https://github.com/open-mmlab/mmselfsup. Our code and pre-trained weights can be obtained from https://github.com/Zhcyoung/BEPH. and can be used to replicate the experiments in this paper. All source code is released under the GNU GPLv3 free software license.

## Supporting information

Supplementary materials, including charts and related word abbreviations that are not presented in the main text

All numerical results of training

## Acknowledgements

The computations in this paper were run on the Siyuan-1 cluster supported by the Center for High Performance Computing at Shanghai Jiao Tong University. Additionally, we are grateful to Ph.D. student Jie Zhou, Ph.D student Xin Yang, Ph.D student Yifan Wang, Ph.D student Yidan Cuifor their useful discussion and the help of other students and teachers in Yu Lab

## Ethical approval

Ethical approval was not required for this study as it did not involve human or animal subjects.

