## Supplementary materials, including charts and related word abbreviations that are not presented in the main text for "A foundation model for generalizable cancer diagnosis and survival prediction from histopathological images"

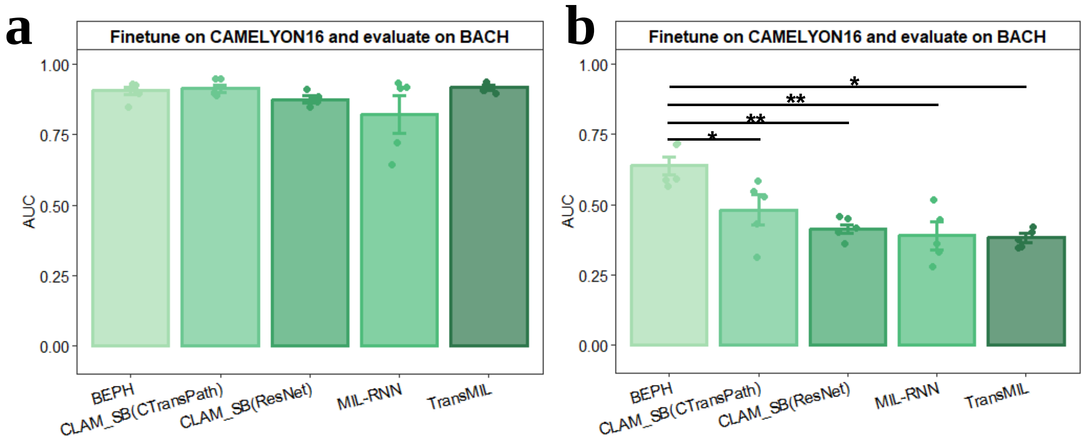


**Supplementary Fig. 1| Performance (AUC) on WSI diagnostic classification. a**, Internal evaluation. models are adapted to the CAMELYON16 dataset and internally evaluated on hold-out validation data. **b**, External evaluation. models are adapted to CAMELYON16 and externally evaluated on another dataset of the same task, BACH. There is no fine-tuning on BACH because there are only 30 available slides

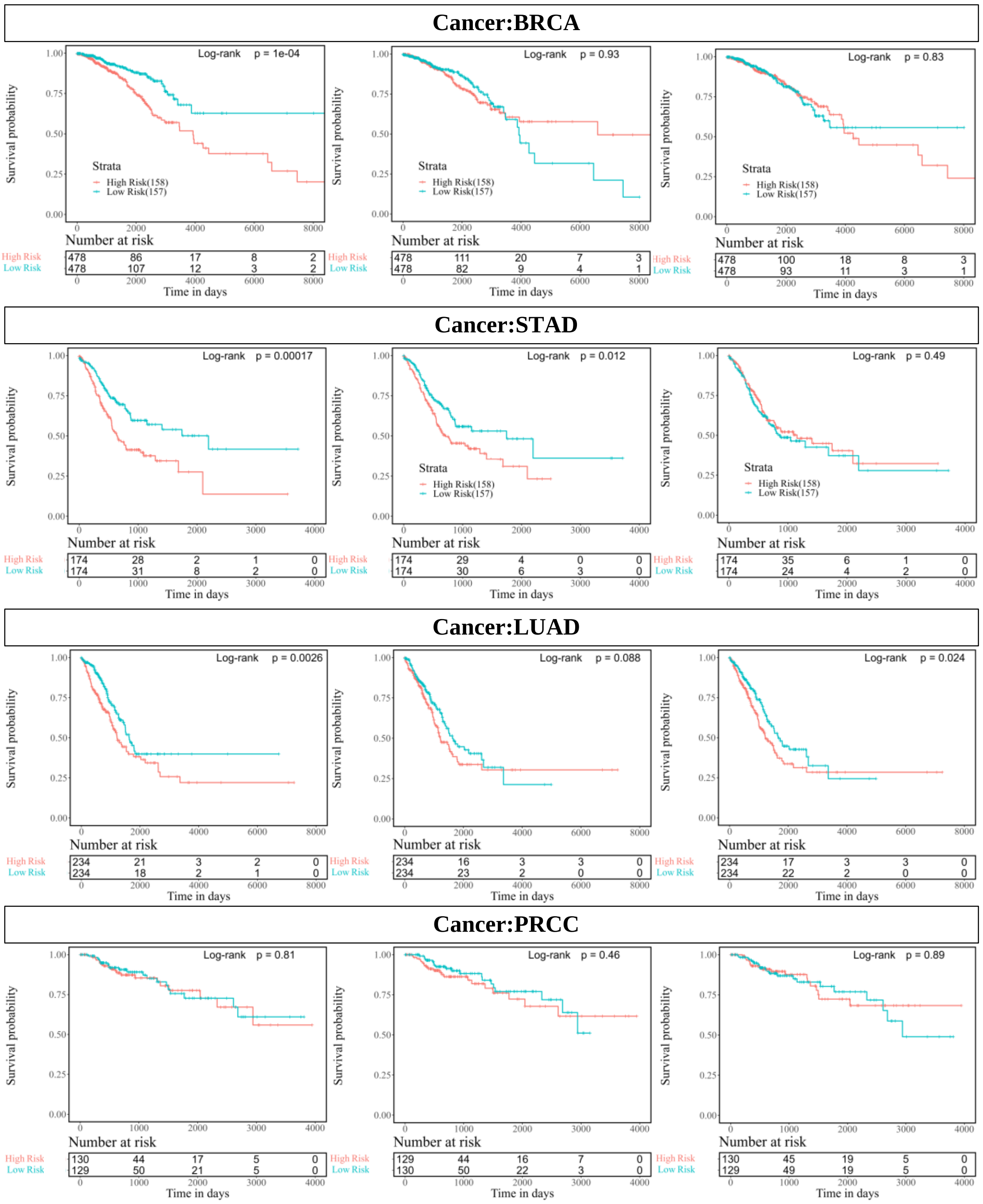


**Supplementary Fig. 2| Survival curves of the four cancers that do not appear in the main body.** On BRCA, STAD, and LUAD, the p-value of BEPH is the smallest and significant. On PRCC, all three methods are not significant.

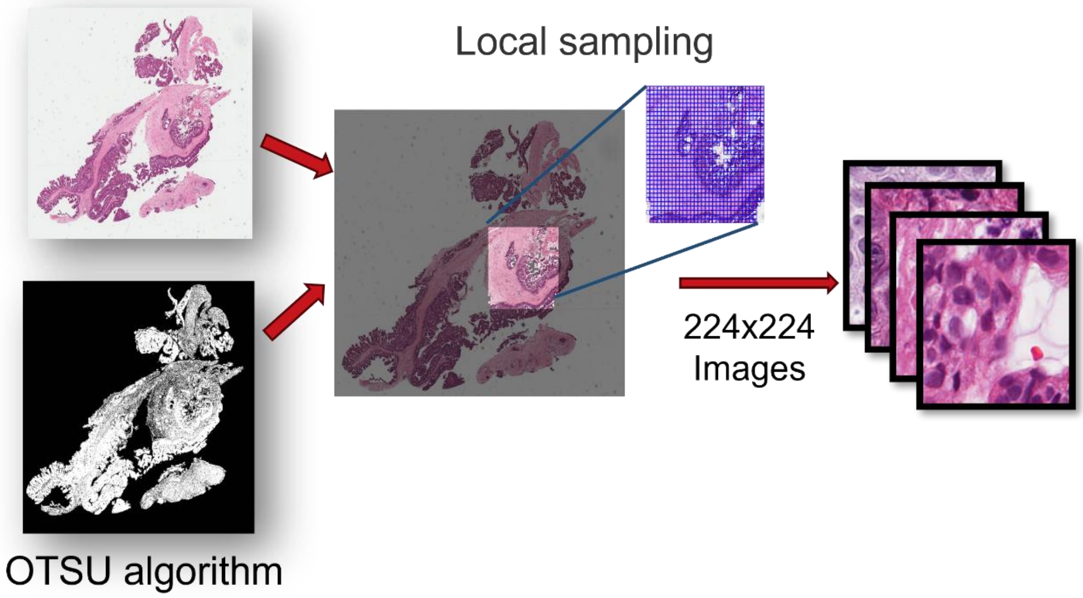


**Supplementary Fig. 3| The details of slicing WSI into unlabeled pre-training datasets.**

**Supplementary Table 1. The correspondence between abbreviations and full names appeared in the article.**

| **Cancer** | **Abbreviations** |
|---|---|
| nonspecific invasive breast cancer | BRCA |
| invasive ductal carcinoma | IDC |
| invasive lobular carcinoma | ILC |
| non-small cell lung cancer | NSCLC |
| lung squamous cell carcinoma | LUSC |
| lung adenocarcinoma | LUAD |
| renal cell carcinoma | RCC |
| colorectal cancer | CRC |
| clear cell renal cell carcinoma | CCRCC |
| chromophobe renal cell carcinoma | ChRCC |
| papillary renal cell carcinoma | PRCC |
| stomach adenocarcinoma | STAD |

**Supplementary Table 2. The backbone of the model on WSI subtyping classification task.**

<table>
<tr><th>Model</th><th>Backbone</th></tr>
<tr><td>CLAM_SB(ResNet)</td><td>the original model from Lu et al, ResNet supervised pretrained on ImageNet-1K</td></tr>
<tr><td>MIL</td><td rowspan="5">DINO pretrained on 256×256 histopathological images</td></tr>
<tr><td>GCN-MIL</td></tr>
<tr><td>DS-MIL</td></tr>
<tr><td>DeepAttnMISL</td></tr>
<tr><td>CLAM_SB(DINO)</td></tr>
<tr><td>HIPT</td><td>DINO pretrained on 256×256 and 4096×4096 pathology images</td></tr>
<tr><td>BEPH</td><td>CLAM_SB whose backbone is BEiTv2 pretrained on 224×224 histopathological images</td></tr>
</table>

**Supplementary Table 3. The backbone of the model on CAMELYON16 and BACH dataset classification task.**

| **Model** | **Backbone** |
|---|---|
| BEPH | CLAM_SB whose backbone is BEiTv2 pretrained on 224×224 histopathological images |
| CLAM_SB(ResNet) | the original model, ResNet supervised pretrained on ImageNet-1K |
| CLAM_SB(CTransPath) | The method from Wang et al. |
| MIL-RCNN | ResNet supervised pretrained on ImageNet-1K |
| TransMIL | SimCLR pretrained on CAMELYON16 |

**Supplementary Table 4. The backbone of the model on WSI survival prediction task**

| Model | Backbone |
|---|---|
| CLAMSurvival (ResNet) | ResNet supervised pretrained on ImageNet-1K |
| ABMIL | DINO pretrained on 256×256 histopathological images |
| GCN-MIL | |
| DS-MIL | |
| DeepAttnMISL | |
| CLAM_SB(DINO) | |
| HIPT | DINO pretrained on 256×256 and 4096×4096 pathology images |
| BEPH | CLAMSurvival whose backbone is BEiTv2 pretrained on 224×224 histopathological images |
